# D^R^e^A^mocracy: A Method to Capitalize on Prior Drug Discovery Efforts to Highlight Candidate Drugs for Repurposing

**DOI:** 10.1101/2023.01.12.523717

**Authors:** Kyriaki Savva, Margarita Zachariou, Marilena Bourdakou, Nikolas Dietis, George M. Spyrou

## Abstract

In the area of drug research, several computational drug repurposing studies have highlighted candidate repurposed drugs, as well as drugs from clinical trial studies in different phases. To our knowledge, the aggregation of the proposed lists of drugs by previous studies has not been extensively exploited towards the generation of a dynamic reference matrix regarding the disease-related frequencies of drug features such as the modes of action, the initial indications and the targeted pathways. To fill this knowledge gap, we performed a weight-modulated majority voting of the modes of action, initial indications and targeted pathways of the drugs in a well-known repository, namely the Drug Repurposing Hub. Our method, D^R^e^A^mocracy, exploits this pile of information and creates frequency tables and finally a disease suitability score for each drug. As a testbed, we applied this method to a group of neurodegenerative diseases (Alzheimer’s, Parkinson’s, Huntington’s and Multiple Sclerosis). A super-reference table with drug suitability scores has been created for all four neurodegenerative diseases and can be queried for any drug candidate against them. Top-scored drugs for Alzheimer’s disease include *agomelatine* and *mirtazapine*, for Parkinson’s disease *apomorphine* and *pramipexole*, for Huntington’s *fluphezine* and *perphezine*, and for Multiple Sclerosis *zonisamide* and *disopyramide*. Overall, D^R^e^A^mocracy is a methodology that leverages the existing experimental and/or computational knowledge (1) to reveal trends in selected tracks of drug discovery research that includes modes of action, targeted pathways and initial indications for the investigated drugs and (2) to score new candidate drugs for repurposing against a selected disease.

## 1. INTRODUCTION

*“…And if you find her poor, Ithaka won’t have fooled you. / Wise as you will have become, so full of experience, / you’ll have understood by then what these Ithakas mean*.*”*^1^ These are the last three verses of the poem “Ithaka” from the Greek poet Constantine Cavafy that deals with the value of the journey in relation to the value of the destination. And this is more relevant than ever in the case of the countless journeys to discover new drugs where the destination is often not what we have expected. Albeit the results of most of the drug discovery trajectories are not successful, the question remains whether we can integrate this knowledge and experience in a democratic way into computational methods that will increase the chances for the next drug discovery efforts.

According to the literature, only a few studies have used this massive amount of knowledge to gain further insight into the drug discovery process. Himmelstein et al., constructed a network that integrated knowledge from a very large number of biomedical studies [1]. Data were integrated from 29 public resources to connect compounds, diseases, genes, anatomies, pathways, biological processes, molecular functions, cellular components, pharmacologic classes, side effects, and symptoms. This approach described more than two million relationships among data points, which could be used to develop models that predict which drugs currently used in the clinic might be best suited to treat any of the 136 common diseases. Moreover, Kropiwnicki et al., (2022) created a tool, named DrugShot (https://maayanlab.cloud/drugshot/), which prioritizes drugs and small molecules associated with biomedical search terms. Apart from listing known associations, DrugShot predicts additional drugs and small molecules related to any search term [2]. In another study, Zhu et al. introduced an approach to knowledge-driven drug repurposing based on a comprehensive drug knowledge graph. They integrated multiple drug knowledge databases in order to develop a drug knowledge graph to predict drug repurposing candidates by using machine learning models [3].

However, to our knowledge, there have been no studies to date that have collectively capitalized on the already available information that has been generated in both computational drug repurposing studies and clinical trials with increased resolution, including modes of action, targeted pathways, and initial indications for the investigated drugs.

Drug repurposing or repositioning, which is the identification of novel uses for existing drugs, has attracted considerable attention during the past few decades as it offers a cost efficient and time effective alternative avenue to therapeutics, compared to *de novo* drug discovery [4], [5]. Due to the growing abundance of different types of omics data in a number of different diseases, numerous drug repurposing approaches have been available and constitute a major part of the drug discovery pipeline for diseases that lack treatment, such as neurodegenerative diseases [6]. In general, traditional drug repurposing attempts to reveal the effect of a drug and its mechanism of action [7], screens available drugs against new targets to reveal novel drug indications [8], investigates drug characteristics such as their structures and side effects [9], or explores the relationships of drugs with diseases [10].

This massive availability of large-scale data, such as microarray gene expression signatures, high-throughput sequencing, or publicly available data in pharmaceutical databases, in combination with high-performance computing has helped accelerate the growth and development of computational drug repurposing approaches. These include data mining, machine learning, and network analysis [11]. Most of the studies in the field of drug repurposing exploit the associations between different biological entities such as drugs, diseases, genes, and adverse drug reactions.

So far, there have been many studies on computational drug repurposing that have attempted to highlight candidate repurposed drugs, as well as clinical trial studies that test drugs in different phases. Both approaches result in long lists of drugs that are usually not exploited further. To our knowledge, this information has not been extensively exploited collectively. We posed the following questions: (1) How can we integrate the available information? (2) How can we leverage the integrated knowledge to generate a knowledge-based model to assist drug discovery?

This available *a priori* knowledge can turn into novel information regarding a specific drug of interest. To do so, we designed and developed a methodology named D^R^e^A^mocracy, which exploits this huge pile of information given by repurposing studies, as well as information available in clinical trials, in a resolution beyond the nominal reference of the drug per study. Specifically, information such as the drugs’ mechanisms of action (MoAs) with their initial indications (Inds) as retrieved from the Drug Repurposing Hub and the targeted pathways (Paths), as obtained from the KEGG Pathway Database, is subjected to weight-modulated majority voting schemes to score and finally prioritise the candidate drugs against a disease.

D^R^e^A^mocracy was applied to four diseases of the neurodegeneration spectrum as a testbed to check the usefulness of this method. Neurodegenerative diseases are a group of diseases that face many challenges in terms of effective availability of treatments, making them a great candidate for drug repurposing. In 2016, it was reported that AD was affecting ∼47 million people, with predictions suggesting this number to reach ∼75 million in 2030. Nichols et al. proposed that dementia would increase from an estimated 57. 4 million cases globally in 2019 to an estimated 152. 8 million cases in 2050 [12]. Moreover, more than 500, 000 Americans have been diagnosed with Parkinson’s Disease (PD), the second most common neurodegenerative disorder [13]. Millions more suffer from other neurodegenerative diseases such as Frontotemporal Dementia (FTD), Huntington’s disease (HD), Amyotrophic Lateral Sclerosis (ALS), Multiple Sclerosis (MS), Spinal Muscular Atrophy (SMA) [14]–[17]; diseases that lack effective treatments today. These diseases carry a great economic burden on public health. In AD, for example, according to the Alzheimer’s Association, the money spent on their patients reached $321 billion in 2022. Alzheimer’s is projected to cost nearly $1 trillion by 2050 [18]. Therefore, there is an urgent need to discover and propose novel drugs for these diseases.

For this reason, we chose two well-studied diseases, Alzheimer’s disease (AD) and Parkinson’s disease (PD), for which there is a lot of information available on repurposing studies and clinical trials, and two diseases with less available respective information, Multiple Sclerosis (MS) and Huntington’s disease (HD).

## 2. MATERIALS AND METHODS

D^R^e^A^mocracy integrates disease-specific drug-lists generated by various drug discovery approaches. The drug collection methods that are selected in this work are: (1) collection of the lists of drugs from the Computational Drug Repurposing Studies (CDRS) and (2) collection of the lists of drugs from the Clinical Trials Studies (CTS). In addition, there is an option to generate a consensus scheme between the data from the two methods. The general pipeline of D^R^e^A^mocracy is presented in Figure 1. In Step 1, the construction of the reference score matrix is carried out, by choosing the method of interest to collect lists of candidate drugs. These lists could be from CDRS, CTS, Experimental Drug Discovery or any other method available that generates drug lists.

**Figure 1:**
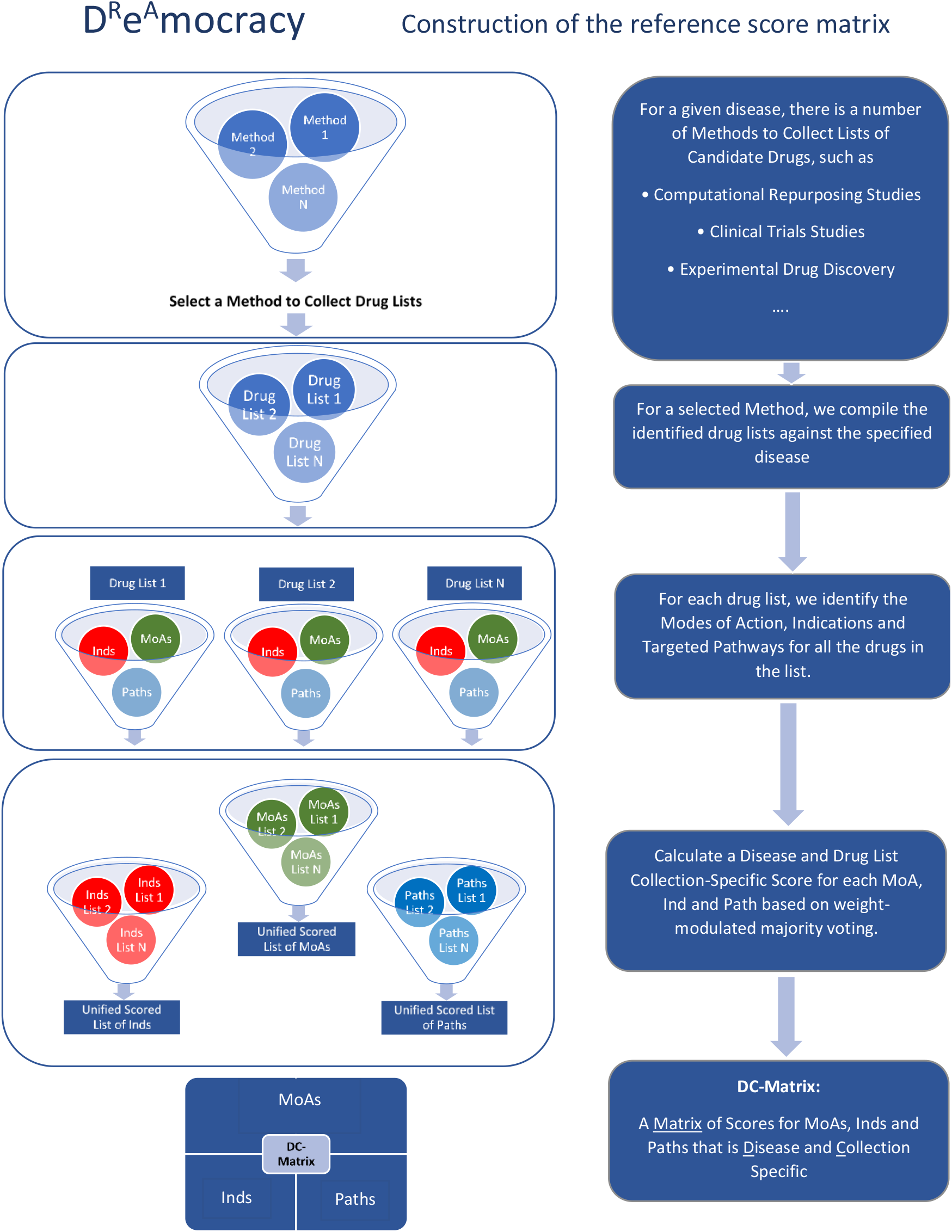
D^R^e^A^mocracy pipeline: Construction of the Reference Score Matrix. Initially, a number of different methods is applied to collect drug lists: computational drug repurposing lists (CDRS; Method 1) on AD, PD, HD and MS, as well as Clinical trials studies (CTS; Method 2) for these diseases. Any method that generates drug lists can also be used (Method N). For each drug list produced by each method, we extracted the mechanisms of action (MoAs), initial indications (Inds) and targeted pathways (Paths) for each drug of the list. We then calculated a specific score for each MoA, Ind and Path (DC score), based on a weight-modulated majority voting. Lastly, a DC matrix that includes the DC scores is generated for each disease and for each method.

The selected diseases to apply this pipeline belong to the spectrum of neurodegeneration as mentioned in the introduction section. For this purpose, we used CDRS on AD, PD, HD and MS, as well as CTS for these diseases. In Step 2, for each drug list for each method, we identified the MoAs, initial Inds and Targeted Paths for all the drugs in the list. In Step 3, we calculated a disease/collection-specific score for each MoA, initial Ind and Path, known as DC score, based on a weighted-modulated majority voting. Lastly, in Step 4, a DC matrix, which is a Matrix of DC Scores for MoAs, Inds and Paths that is disease and method specific, is generated.

The second part of the pipeline (Figure 2) includes the assignment of a disease/collection-specific score to a drug, based on the DC-Matrices of D^R^e^A^mocracy. For any drug or a set of drugs of interest, one can query the D^R^e^A^mocracy-generated reference table to obtain the respective Total Composite Score of the drug based on the disease of interest.

**Figure 2:**
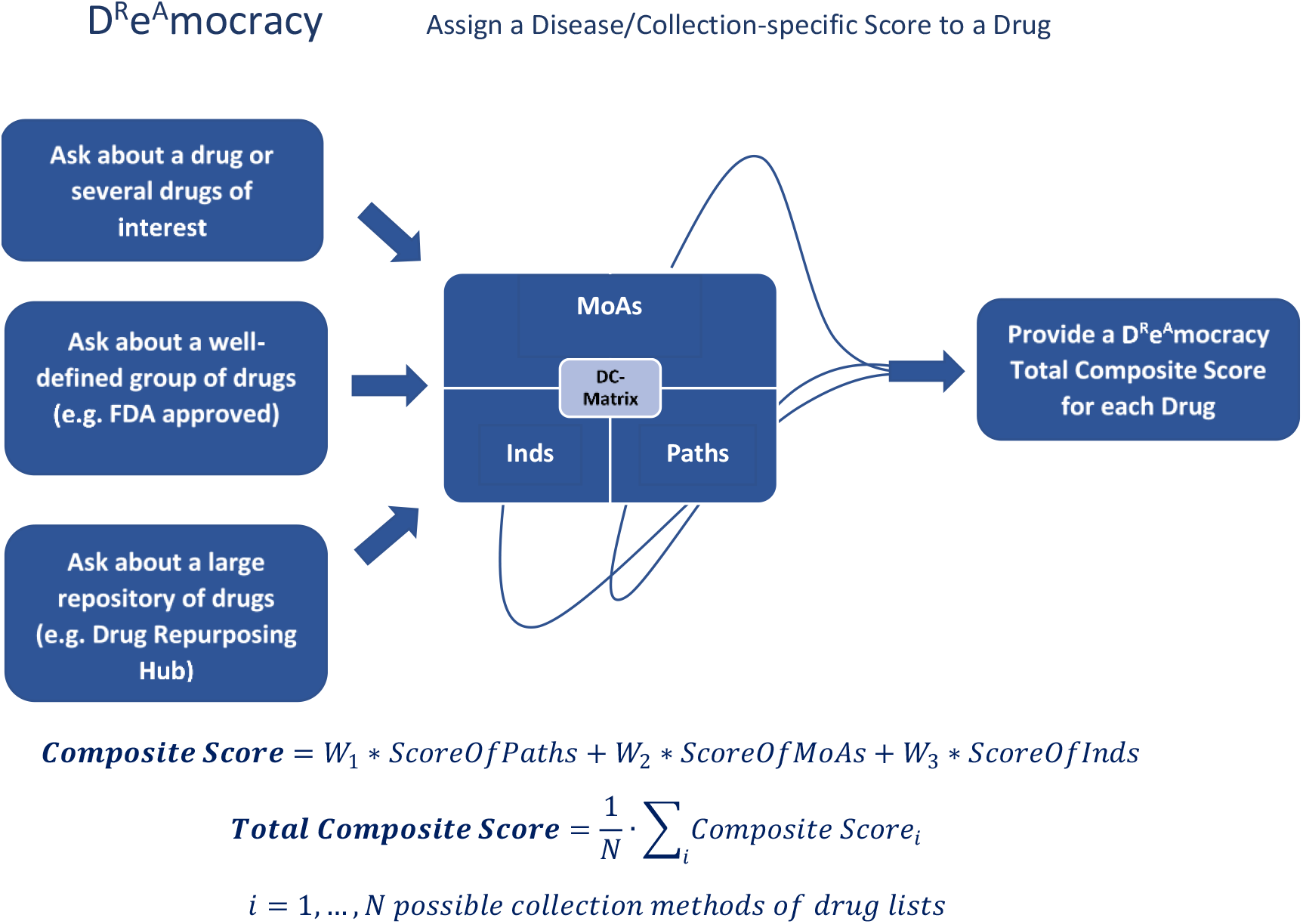
D^R^e^A^mocracy pipeline: Assigning a Disease/Collection-specific Score to a drug. We assign a disease/ collection-specific score for each Drug based on the DC-Matrices of D^R^e^A^mocracy. A Composite score is then generated to combine scores for MoAs, Paths and initial Inds for both CTS and CDRS lists. A final total Composite score for the Composite CTS and Composite CDRS score is generated.

### 2.1 Data Selection

For the CDRS collection method, initial data were retrieved from PubMed using the query type: (repurpos*[tiab] OR reposition*[tiab]) AND (the disease of interest e.g., Alzheimer*[tiab]). The output of the query was filtered to keep only research papers describing computational drug repurposing results for the each of the selected diseases published in the past 10 years (Supplementary Tables 1, 2, 3 and 4).

For the CTS collection method, Clinical Trials information was downloaded from www.clinicaltrials.gov as of 24^th^ of January 2022 for each disease separately. As a keyword, the name of the disease was used in the query. The resulting text tables were downloaded and manually curated to keep only studies whose status was either “Completed”, “Actively Recruiting”, “Recruiting”, “Not yet recruiting” and “Active, not recruiting”. Moreover, in the interventions tab, only drugs were kept. Unique drugs were saved, by keeping the clinical trial that was in the highest phase if multiple studies were available for the same drug.

### 2.2. Construction of the Reference Score Matrix (DC-Matrix)

The drug collection for each disease (AD, PD, MS, HD) and for each method (CDRS, CTS) was searched within the Drug Repurposing Hub Database (https://clue.io/repurposing). Drug indications, modes of action and targets were extracted for each drug in each list. Using the drug target information from the database, we mapped the pathways that drug targets are involved using EnrichR R package [19]. We collected all pathways that the drug targets were members. Then, for each disease, a table of ranked lists for (a) Indications (Inds) (b) Modes of Action (MoAs) and (c) Pathways (Paths) per disease were generated from (1) the CDRS method and (2) the CTS method. These reference tables, also known as DC-Matrices (Drug List Collection Matrices), were prepared for each of the four neurodegenerative diseases. The frequency (Freq) of each drug information (Ind, MoA, Path) as well as the number of initial studies (ListCount) introducing this drug information were calculated. Both metrics, Freq and ListCount, were then normalized in the unit interval. The Lists were ranked based on the normalized frequency (Norm_Freq) and the normalized count of lists (Norm_ListCount) according to the following equation:

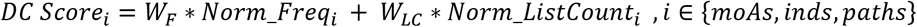

Where W_F_ and W_LC_ are the weights of importance of the normalized frequency and normalized count of lists for the MoAs, Inds and Paths. In this first application of the method, we set W_F_ = 0. 4 and W_LC_=0. 6. A higher weight was given to the ListCount because of the assumption that detection of a signature in as many drug lists as possible, is more important than the frequency of the signature by a single study/drug list.

Finally, to avoid random insertions in the DC-Matrix Lists, these scored lists were filtered using a hypergeometric test. Then we kept only the MoAs, Inds and Paths with a hypergeometric test p-value lower than 0. 05.

### 2.3. Assign a Disease/Collection-specific Score to a Drug

Figure 2 describes the case where we can ask D^R^e^A^mocracy about (1) a drug or several drugs of interest, (2) a well-defined group of drugs (e. g. FDA approved), (3) a large repository of drugs (e. g. Drug Repurposing Hub). To proceed with the drug(s) scoring against a selected disease, we extract the corresponding Inds, MoAs and Paths of the drug(s) and we combine the corresponding MoAs, Paths and Inds scores from the DC-Matrix into a Composite score for each drug against the selected disease using the following equation:

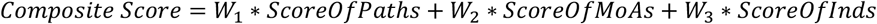

In this first application of the method, we set W_1_ = 0. 4, W_2_ = 0. 4 and W_3_=0. 2. Paths and MoAs were given equal and higher weights than the initial Inds since the first two give an actual description of the features of the drug rather than the initial use of the drug.

In the case of combination of all the drug collection methods, then a Total Composite Score can be calculated as an average of the Composite Scores for each data collection method (e. g. CDRS and CTS) as shown in the following equation:

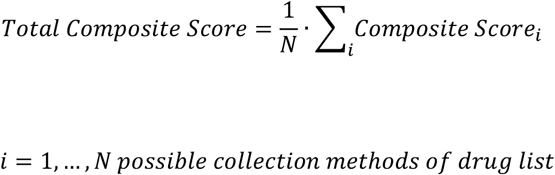

### 2.4. The whole Drug Repurposing Hub through the looking glass of D^R^e^A^mocracy

We performed a large-scale computation by calculating the *Total Composite Score* for each drug in the Drug Repurposing Hub with a total of 4421 drugs for four diseases: AD, PD, HD and MS. The whole pipeline of D^R^e^A^mocracy however, can be applied in multiple diseases given sufficient availability of drug-related a-priori information. This can be extended to a greater number of diseases as shown in Figure 3 (*N* diseases) and also, any other database with the same type of can be used. Moreover, in addition to the different diseases that can be tested, different methods to collect drug lists can also be used, such as experimental drug discovery studies. Notably, a better Composite score can be given when more data are available for the disease of interest, as a higher diversity can be added to the methodology. Therefore, to obtain more reliable results, it is necessary to create a minimum of three drug lists per disease to proceed with the creation of its reference table.

**Figure 3:**
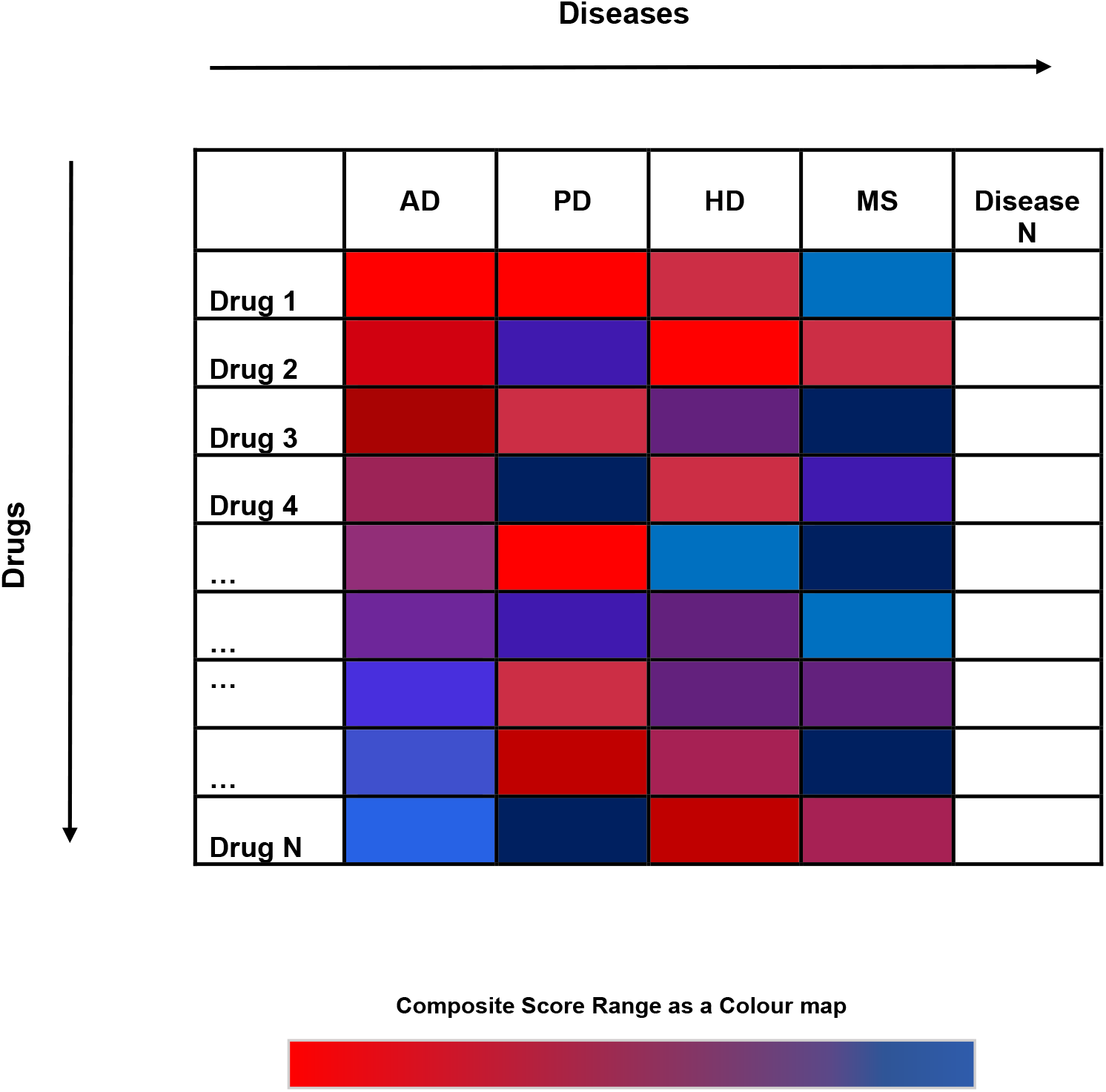
Schematic for a large-scale application of D^R^e^A^mocracy for various diseases. By following the D^R^e^A^mocracy pipeline, the total Composite scores can be calculated for any disease of interest that has available prior knowledge, either on Clinical Trial Studies or on Computational Drug repurposing Studies. Drug Repurposing Hub was used in this work to extract the available information regarding the drugs. In the current scheme, the drugs in the table are considered sorted based on their score in the first column (AD).

## 3. RESULTS

### 3.1. Reference tables for each disease, type of analysis and drug feature

As previously discussed, lists of candidate repurposed drugs from computational drug repurposing studies, as well as clinical trial studies for four neurodegenerative diseases (AD, PD, HD, MS), were collected to utilise this integrated information available for a better and more effective suggestion of repurposed drugs for a given disease.

By calculating the DC-score for the aggregated MoAs, Paths and initial Inds, we detected the top-ranking ones per disease and drug collection method. The DC-Matrix for AD CDRS list, revealed as top scoring drug features the *serotonergic synapse, serotonin receptor antagonist* and *hypertension* for Paths, MoAs and initial Inds respectively. *Serotonin receptor antagonism* was also found in the top four MoAs for PD CDRS and CTS DC-Matrices, with a DC score of 0.56 and 0.75 respectively. For the initial Inds, *hypertension* was detected as not only the highest score for AD, but also for the PD CDRS DC-Matrix table (Figure 4). There are many studies and meta-analyses available that show the association of antihypertensives and AD, and specifically the effect that they have in decreasing the risk of AD [20]–[22]. Moreover, antihypertensives, such as centrally-acting dihydropyridine use and high cumulative doses of ACEIs, were shown to have a possible association with reduced PD incidence in hypertensive PD patients [23].

**Figure 4:**
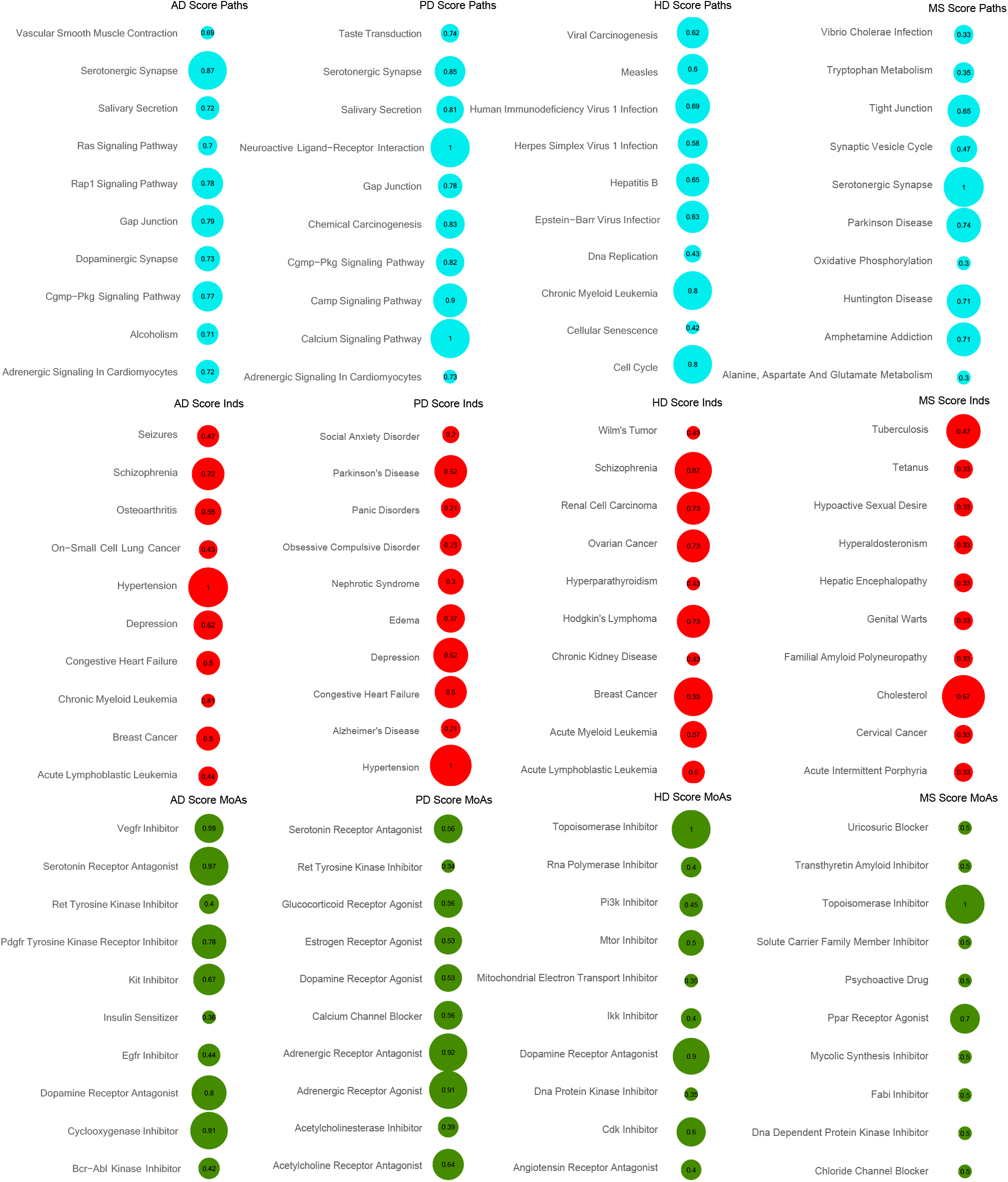
DC-Matrix for CDRS. DC-matrices were generated for each disease per signature (Paths, MoAs and Inds) using the CDRS lists. The top 10 scored features are shown for each disease in a bubble grid chart. The number in the bubbles indicates the DC score of each modality (also encoded as the bubble size). Blue colour depicts Paths, red colour initial Inds and green colour MoAs.

The *cAMP signalling pathway*, which was identified as the highest scoring pathway in HD CTS DC-Matrix tables (with a score of 1) and in the top three of PD CDRS and CTS (0.9 and 0.79 respectively), has shown to play an important role in mediating neurotransmitters and regulating numerous cell functions, such as synaptic plasticity in neurons. In PD, dysregulated *cAMP signalling pathway* is associated with levodopa-induced dyskinesia, whereas its dysfunction was also found in HD. [24], [25].

Moreover, *topoisomerase inhibitor* was detected as the highest scoring MoA in both HD and MS CDRS reference tables with a score of 1 in both diseases. The DNA *topoisomerase inhibitor* mitoxantrone is a drug used for patients with worsening relapsing-remitting and secondary progressive multiple sclerosis and hepatocellular carcinoma [26], [27]. On the other hand, in the CTS DC-matrix *glutamate receptor antagonist* and *sodium channel blocker* were detected as the top ones respectively (Figure 5). As shown by Abd-Elrahman *et al*. (2017), blocking glutamate receptors and specifically the metabotropic glutamate receptor 5 (mGluR5) in a HD mouse model, an improvement in cellular, motor, and cognitive skills was observed. In addition, the *sodium channel blocker safinamide*, which was tested in phase III trials for PD, could also be used for MS to protect against axonal degeneration regarding to Morsali and colleagues, as it preserved the integrity and function of the axon in two rat models of MS [29]. Full lists of DC-matrices for CDRS and CTS per disease are available in Supplementary Tables 8-15.

**Figure 5:**
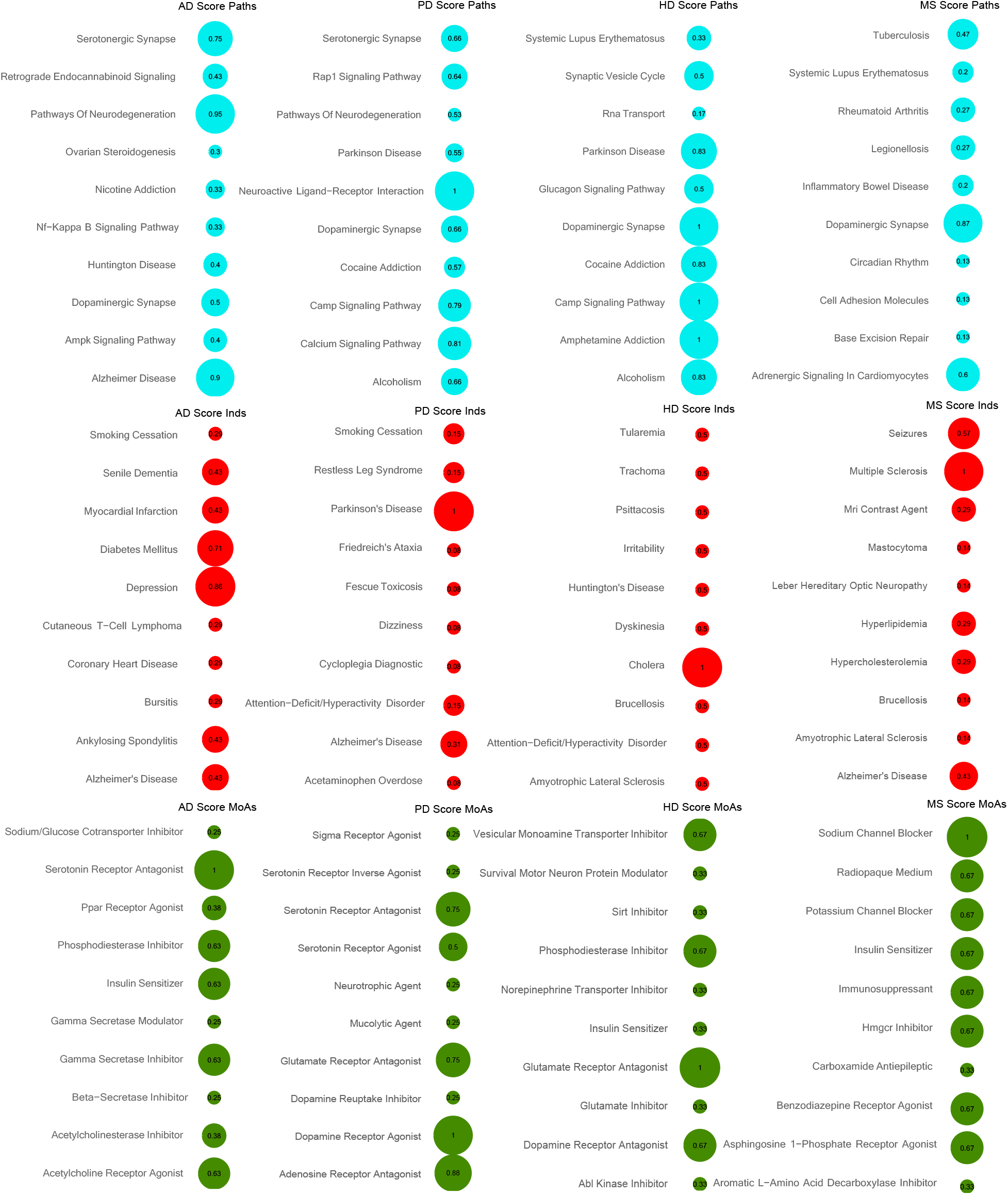
DC-Matrix for CTS. DC-matrices were generated for each disease per signature (Paths, MoAs and Inds) using the CTS lists. The top 10 scored modalities are shown for each disease in a bubble grid chart. The number in the bubbles indicates the DC score of each modality (also encoded as the bubble size). Blue colour depicts Paths, red colour initial Inds and green colour MoAs.

### 3.2. Common signatures in CTS vs CDRS for neurodegenerative diseases

As a next step, we compared the CDRS and CTS signatures (MoAs, Inds and Paths) detected for each disease independently. The top signatures were selected using a hypergeometric test and a p-value of <0.05, as shown in Figure 6. For AD, commonalities in CDRS and CTS were found across all components of the signature. For MoAs, six commonalities were found, some of which are highly associated with AD. These include the *serotonin receptor antagonist, acetylcholinesterase inhibitor* and *HMGCR inhibitors*. All three MoAs have been associated with neurodegeneration and specifically AD [30]–[32]. Moreover, at present, *acetylcholinesterase inhibitors*, such as donepezil and rivastigmine, are the main class of drugs currently used for the treatment of AD [32].

**Figure 6:**
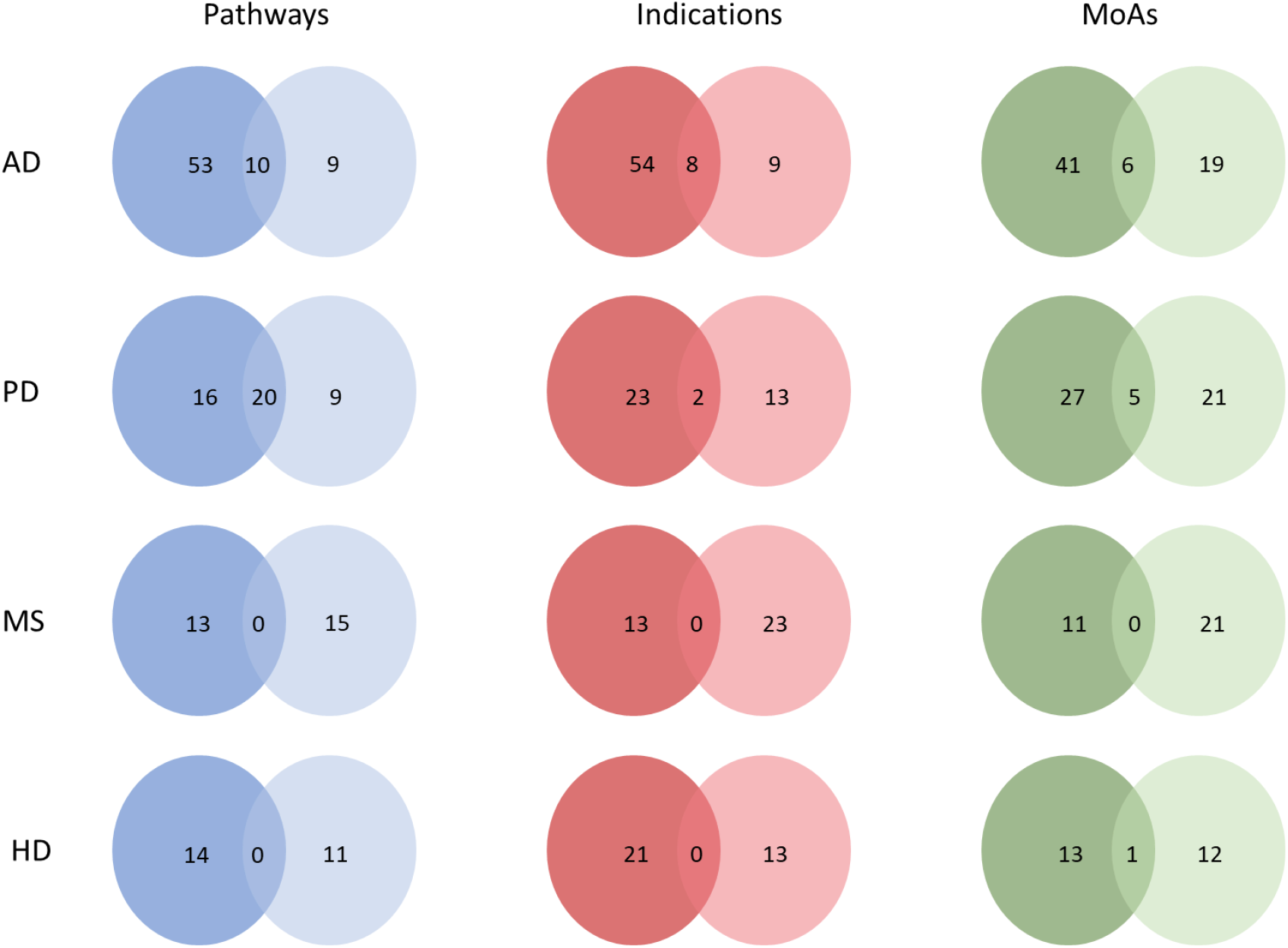
Signature comparison across different diseases. CDRS represent signatures detected from drugs in computational drug repurposing lists and CTS represent signatures detected from drugs in current clinical trials for each disease respectively. CDRS lists are depicted using the darker colour and CTS lists with a lighter colour for each property of the drugs (Pathways, Indications and MoAs).

For initial Inds, eight commonalities were detected. These include cutaneous *T-cell lymphoma (CTCL), depression, heart attack, ankylosing spondylitis, bursitis, coronary heart disease, exercise-induced bronchoconstriction (EIB)*, and *myocardial infarction*. Regarding *CTCL, HDAC inhibitors*, such as romidepsin and vorinostat, are used for treatment, and have been reported to exhibit neuroprotective actions, something that could be achieved by multiple MoAs, either through the suppression of pro-apoptotic factors or through the release of pro-inflammatory factors of activated microglia [33], [34]. Furthermore, *depression* has been very common to patients with AD, and also, a history of *depression* may increase risk for developing AD later in life [35]. Additionally, ten common Paths were found, *serotonergic synapse, AMPK signalling pathway, dopaminergic synapse, viral myocarditis, ovarian steroidogenesis, glycerophospholipid metabolism, PPAR signalling pathway, circadian rhythm, arachidonic acid metabolism* and *notch signalling pathway*.

For PD, commonalities in CDRS and CTS were found across all components of the signature, as shown for AD. For MoAs, five commonalities were detected such as *dopamine receptor agonist, serotonin receptor antagonist, dopamine precursor, aromatic L-amino acid decarboxylase inhibitor* and *norepinephrine precursor*. Dopamine receptor agonists are currently tested for PD, and some of them have been approved by the FDA for the disease [36].

For the initial Inds of PD, two commonalities were found between CDRS and CTS of PD and AD. Furthermore, 20 common Paths were identified such as the *dopaminergic synapse, serotonergic synapse, cAMP signalling pathway, neuroactive ligand-receptor interaction, calcium signalling pathway* and others, some of which will be discussed in the next sections.

For HD, commonalities in CDRS and CTS were found only in MoAs. Specifically, one common element was detected, the *dopamine receptor antagonist*. One of the two FDA approved drugs for HD, known as tetrabenazine, depletes dopamine by inhibiting the vesicular monoamine transporter 2 (VMAT2) [37]. On the other hand, for MS, no commonality was detected in either signature.

These commonalities across CDRS and CTS suggest that repurposing studies and clinical trials agree in some aspects, however, there are also many unique findings.

Moving on from the detection of common signatures in CDRS and CTS for the neurodegenerative diseases under investigation, we wanted to explore whether frequencies of the features of the drugs are consistent in the aforementioned lists. Hence, we performed a comparison across diseases to detect common signatures (MoAs, Inds and Paths). As it can be observed in Figure 6, for MS and HD, no commonalities were detected, therefore, frequency comparison was performed only for AD and PD (Figure 7). For this comparison, only statistically significant features of drugs were used (hypergeometric test and a p-value <0.05). For AD Path comparison, we found a significant relationship using regression analysis (p-val= 4.751e-07) between frequencies of CDRS and CTS lists, with a r^2^ of 0.9638± 0.071. R squared (r2) is a statistical test that shows how well the data fit the regression model (the goodness of fit).

**Figure 7:**
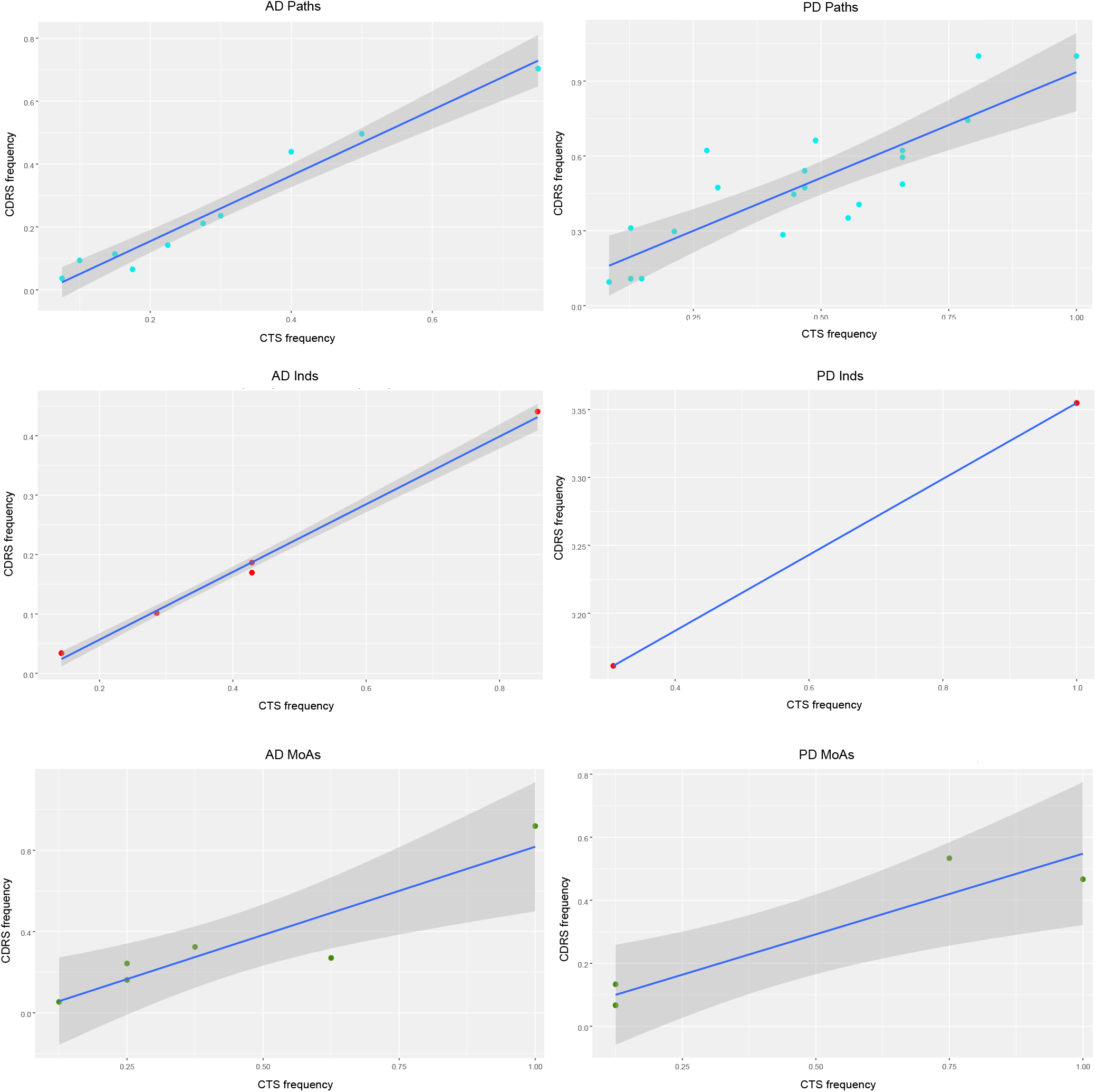
Frequency comparison in AD and PD. Points represent the normalised frequency of signatures in CDRS (y-axis) and CTS (x-axis). Linear regression is used to model the relationship between the two variables and estimate the value of a response by using a line-of-best-fit. Blue colour dots depict Paths, red colour dots Inds and green colour dots MoAs.

For instance, frequency comparison, *serotonergic synapse, dopaminergic synapse* and *AMPK signalling pathway* are the top scored Paths detected that have analogous frequencies in both CDRS and CTS. For AD initial Inds, we found a significant relationship (p-val= 4.535e-08) between frequencies of CDRS and CTS lists, with a r^2^ of 0.9947±0.017. For instance, *depression* is the highest scored pathway, with a normalised frequency score 0.85 in CTS and 0.45 in CDRS. In CDRS list, *depression* scored third, however, it is the top scored common MoA with CTS. Moreover, for the comparison of AD MoAs, we found a significant relationship (p-val= 0.008) between frequencies of CDRS and CTS lists, with a r^2^ of 0.8533±0.18. For instance, *serotonin receptor antagonist* was found to be the top scored MoA, with an analogous frequency in both CDRS and CTS (0.92 and 1 respectively).

For comparison of the PD Paths, we found a significant relationship (p-val= 2.701e-06) between frequencies of CDRS and CTS lists, with an r^2^ of 0.7146± 0.13. For instance, *calcium signalling pathway* and *neuroactive ligand-receptor interaction* were detected as the top scored Paths, with the former having a normalised frequency of 1 and 0.81 for CDRS and CTS respectively, and the latter having a normalised frequency of 1 for both CDRS and CTS. For PD initial Inds, *Alzheimer’s disease* scored low in both CDRS and CTS (0.2 and 0.3 respectively). Lastly, for PD MoAs, we found a significant relationship (p-val= 0.01593) between frequencies of CDRS and CTS lists, with a r^2^ of 0.8904± 0.10. For example, *Dopamine receptor agonist* had a top score for CTS (score 1) whereas for CDRS had a score of 0.47. Moreover, *serotonin receptor antagonist* had a score of 0.57 for CDRS and 0.75 for CTS. These results show that for the common signatures between CDRS and CTS lists have a consistent ranking in both lists.

### 3.3. Commonalities and differences in signatures across neurodegenerative diseases

For the next part of the analysis, we wanted to find commonalities across the four neurodegenerative diseases under investigation, in order to detect common patterns that could be possibly observed behind neurodegeneration.

A disease-to-disease network was created for both CDRS and CTS lists results to show commonalities between diseases, as well as commonalities and differences between CTS and CDRS lists (Figure 8). In CTS, results are more consistent and all components of the signature (MoAs, Paths and Inds) are shared by all four neurodegenerative diseases. On the other hand, for CDRS lists, for the two diseases (AD and PD) for which the data was more abundant, we observed a stronger consistency in all components of the signature. Connections with MS and HD are weaker, with AD and HD sharing only common Inds, HD and MS MoAs and Inds, MS and PD Paths and MoAs whereas PD and HD have zero commonalities.

**Figure 8:**
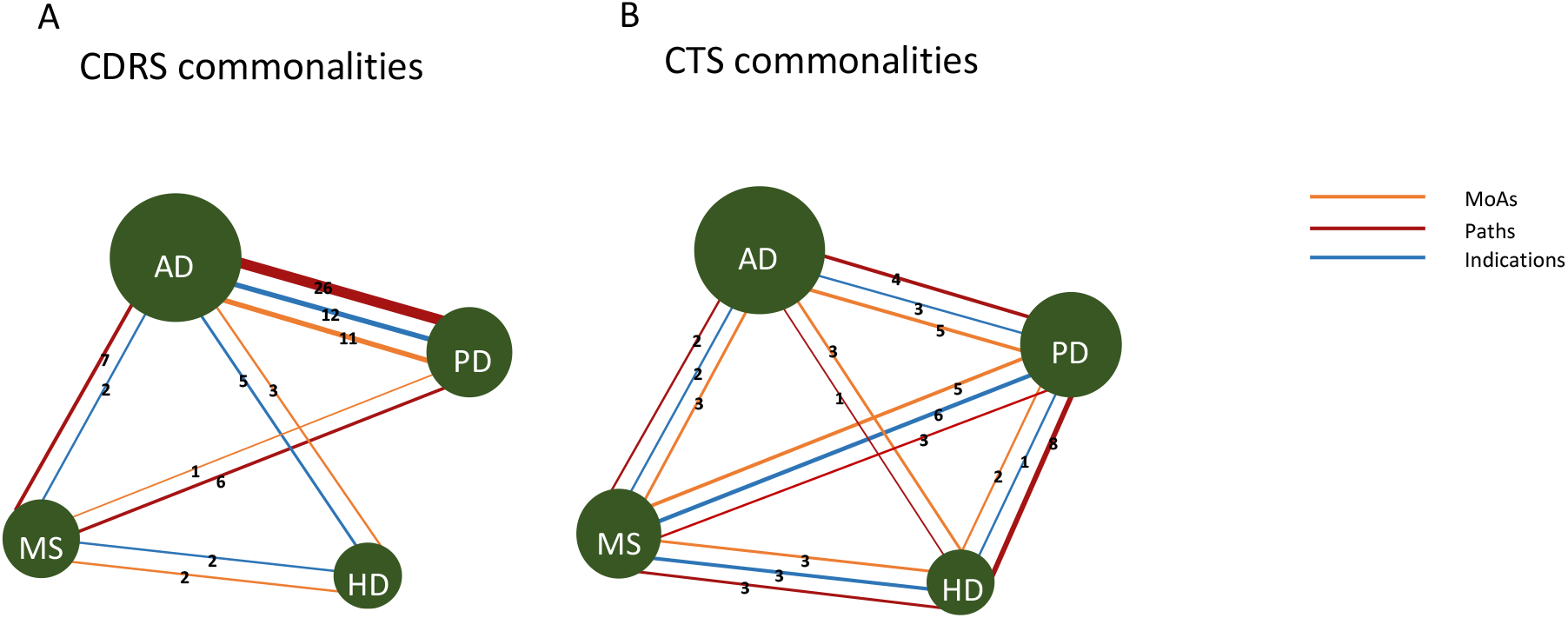
Commonalities across the four neurodegenerative diseases. Disease-disease network. Nodes are the four neurodegenerative diseases and edges are MoAs, Paths and Inds. **A**. Network of CDRS commonalities **B**. Network of commonalities of CTS. Edges indicate the number of MoAs, Paths and Inds that are shared between diseases. The size of the nodes indicates the number of drugs available for each disease.

More specifically, for disease comparisons for initial Inds using the CDRS lists, commonalities were only found pairwise. For example, for AD and HD, *acute lymphoblastic leukemia (ALL), and schizophrenia* were detected, among others. One of the drugs that is used to treat leukemia in adults, is amsacrine, a *topoisomerase inhibitor*. As we discussed in the previous section, mitoxantrone, which is also a *topoisomerase inhibitor*, is used for worsening relapsing-remitting and secondary progressive multiple sclerosis [38]. Hence, these results could indicate a connection between ALL and neurodegenerative diseases.

For AD and PD, *depression, hypertension* and others were detected. For MoAs, commonalities were found both in pairwise and among 3 diseases. The *serotonin reuptake inhibitor* was detected as the common MoA among AD, PD, and MS. Drugs in this category are used in neurology/psychiatry, such as the anti-depressants paroxetine, sertraline, trazodone and others. For AD and HD, *angiotensin receptor antagonist* and *dopamine receptor antagonist* were detected. For AD and PD, 10 common MoAs, such as *acetylcholinesterase inhibitor, dopamine receptor agonist*, and *serotonin receptor antagonist* were identified. The cholinergic system, whose main neurotransmitter is acetylcholine, which is broken down by the acetylcholinesterase enzyme, is often associated with neurodegenerative diseases and hence, inhibitors of this enzyme are frequently used for the treatment of these diseases [36].

For disease comparisons for Paths, commonalities were found both pairwise and among 3 diseases. For AD, PD and MS, *amphetamine addiction, serotonergic synapse* and others were detected. For AD and MS, *ABC transporters, tight junction*, and for PD and MS, *phenylalanine metabolism* were detected. *ABC transporters pathway* is targeted by drugs such as repaglinide, glyburide, which are used in diabetes mellitus and hyperglycaemia. There is biological and epidemiological evidence to support the idea that people with type II diabetes are at increased risk of developing all types of dementia [39]. For AD and PD, a total of 22 Paths were detected; *AMPK signalling pathway, calcium signalling pathway and dopaminergic synapse*, were detected among others. AMPK activation has been shown to play a preventive role in AD, as many studies have shown [40]. On the other hand, other studies have reported that AMPK activation has an aggravating effect on the development of AD [41]. These findings make the therapeutic potential of AMPK in AD controversial.

For disease comparisons for initial Inds using the CTS lists, commonalities were found both pairwise as well as across diseases. *Alzheimer’s disease* was found as a common indication for all four neurodegenerative diseases. In addition, *mastocytoma* was found to be common in AD and MS. *Friedreich’s ataxia*, among others, was found to be common in PD and MS. *Senile dementia* was found to be common in AD and PD, whereas *amyotrophic lateral sclerosis (ALS)* and *hypercholesterolemia* were found to be common in MS and HD, among others. The *norepinephrine transporter inhibitor* was found to be a common MoA in the four neurodegenerative diseases. Changes in norepinephrine reuptake, which is carried out by the norepinephrine transporter, are observed in many neurodegenerative diseases, such as AD and PD [42], [43].

Moreover, *glutamate release inhibitor* and *serotonin receptor antagonist* were found to be common in AD and PD, among others. The *glutamate inhibitor* was found to be common in HD and MS, while the *calcium channel modulator* was found to be common in PD and MS, among others. Alterations in calcium channels have been implicated in several neurodegenerative diseases, such as AD, PD and HD [44]. For disease comparisons for Paths, *dopaminergic synapse* was found as a common indication in all four neurodegenerative diseases. Alterations in the dopaminergic system are more common in PD, where a large proportion of dopaminergic neurons in the Substantia Nigra pars compacta are lost, but are also frequent in AD [45], [46]. The complete lists of comparisons can be found in the Supplementary Tables 5 and 6.

### 3.4. Generation of the super-reference table of drugs

Once the individual tables for each signature were created, the next step was to generate a single reference table for each disease, including a Total Composite Score for all components of the signature (MoAs, Paths and Inds) for both collection methods. A snapshot of the super-reference table generated using the drug library of drug repurposing hub is shown in Table 2. The highest 100 scoring drugs for each disease were selected using the total Composite score.

**Table 1:** Normalised frequency of signatures in CDRS and CTS of AD and PD as shown in Figure 7. Common signatures between CTS and CDRS are shown for each of the two diseases, along with their normalised frequency score.

| **Paths AD** | **CTS** | **CDRS** |
| --- | --- | --- |
| Serotonergic synapse | 0.75 | 0.70 |
| Dopaminergic synapse | 0.50 | 0.50 |
| AMPK signaling pathway | 0.40 | 0.44 |
| Ovarian steroidogenesis | 0.30 | 0.24 |
| Arachidonic acid metabolism | 0.28 | 0.21 |
| PPAR signaling pathway | 0.23 | 0.14 |
| Glycerophospholipid metabolism | 0.15 | 0.11 |
| Viral myocarditis | 0.10 | 0.09 |
| Notch signaling pathway | 0.18 | 0.07 |
| Circadian rhythm | 0.08 | 0.04 |
| **Inds AD** | **CTS** | **CDRS** |
| Depression | 0.86 | 0.44 |
| Myocardial infarction | 0.43 | 0.19 |
| Ankylosing spondylitis | 0.43 | 0.17 |
| Bursitis | 0.29 | 0.10 |
| Coronary heart disease | 0.29 | 0.10 |
| Cutaneous T-cell lymphoma (CTCL) | 0.29 | 0.10 |
| Exercise-induced bronchoconstriction (EIB) | 0.14 | 0.03 |
| Heart attack | 0.14 | 0.03 |
| **Moas AD** | **CTS** | **CDRS** |
| Serotonin receptor antagonist | 1.00 | 0.92 |
| Acetylcholinesterase inhibitor | 0.38 | 0.32 |
| Insulin sensitizer | 0.63 | 0.27 |
| HMGCR inhibitor | 0.25 | 0.24 |
| Angiotensin receptor antagonist | 0.25 | 0.16 |
| Lipoprotein lipase activator | 0.13 | 0.05 |
| **Paths PD** | **CTS** | **CDRS** |
| Neuroactive ligand-receptor interaction | 1.00 | 1.00 |
| Calcium signaling pathway | 0.81 | 1.00 |
| Camp signaling pathway | 0.79 | 0.74 |
| Cgmp-PKG signaling pathway | 0.49 | 0.66 |
| Serotonergic synapse | 0.66 | 0.62 |
| Salivary secretion | 0.28 | 0.62 |
| Dopaminergic synapse | 0.66 | 0.59 |
| Gap junction | 0.47 | 0.54 |
| Alcoholism | 0.66 | 0.49 |
| Amphetamine addiction | 0.47 | 0.47 |
| Vascular smooth muscle contraction | 0.30 | 0.47 |
| Taste transduction | 0.45 | 0.45 |
| Cocaine addiction | 0.57 | 0.41 |
| Parkinson disease | 0.55 | 0.35 |
| Synaptic vesicle cycle | 0.13 | 0.31 |
| Pancreatic secretion | 0.21 | 0.30 |
| Morphine addiction | 0.43 | 0.28 |
| Systemic lupus erythematosus | 0.15 | 0.11 |
| Tyrosine metabolism | 0.13 | 0.11 |
| Phenylalanine metabolism | 0.09 | 0.09 |
| **Inds PD** | **CTS** | **CDRS** |
| Parkinson's disease | 1.00 | 0.35 |
| Alzheimer's disease | 0.31 | 0.16 |
| **Moas PD** | **CTS** | **CDRS** |
| Serotonin receptor antagonist | 0.75 | 0.53 |
| Dopamine receptor agonist | 1.00 | 0.47 |
| Dopamine precursor | 0.13 | 0.13 |
| Aromatic L-amino acid decarboxylase inhibitor | 0.13 | 0.07 |
| Norepinephrine precursor | 0.13 | 0.07 |

**Table 2:**
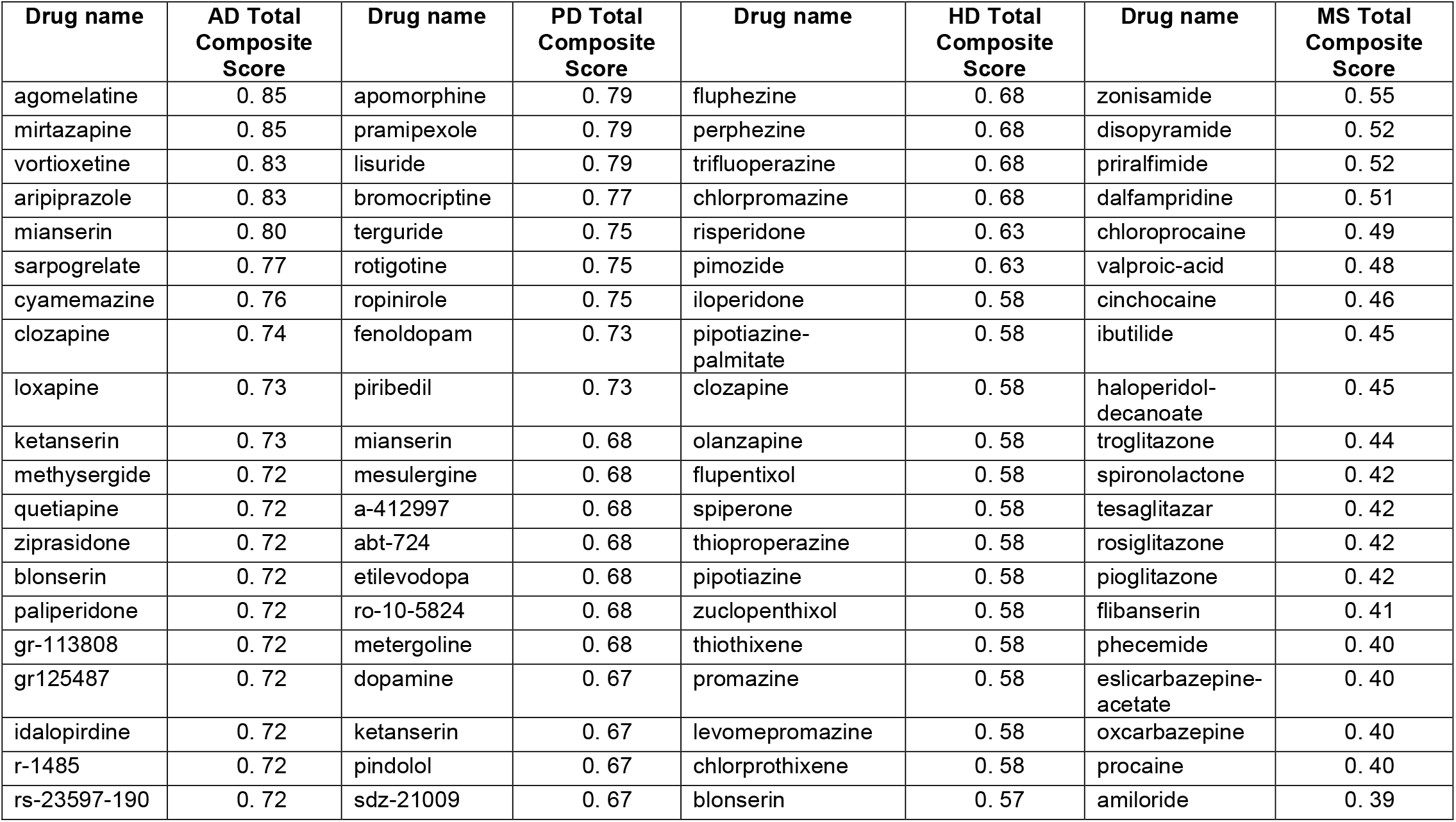
Super-reference table. A snapshot of the super-reference table (top 20 drugs ranked by each disease’s total Composite score separately). In this sub-table (see Supplementary Table 7 for whole table) the final total Composite score is shown for all four neurodegenerative diseases, using the Drug Repurposing Hub drugs as input to D^R^e^A^mocracy.

From the CDRS Composite scores, common drugs across diseases were found in pairwise. Three common drugs were detected between AD and PD (ketanserin, agomelatine and mirtazapine). For AD and HD, 38 common drugs were found, such as iloperidone, olanzapine, amisulpride, and risperidone. For HD and MS, 11 common drugs were found in the top 100 drugs for both diseases including idarubicin, daunorubicin, amsacrine and teniposide (Figure 9B).

**Figure 9:**
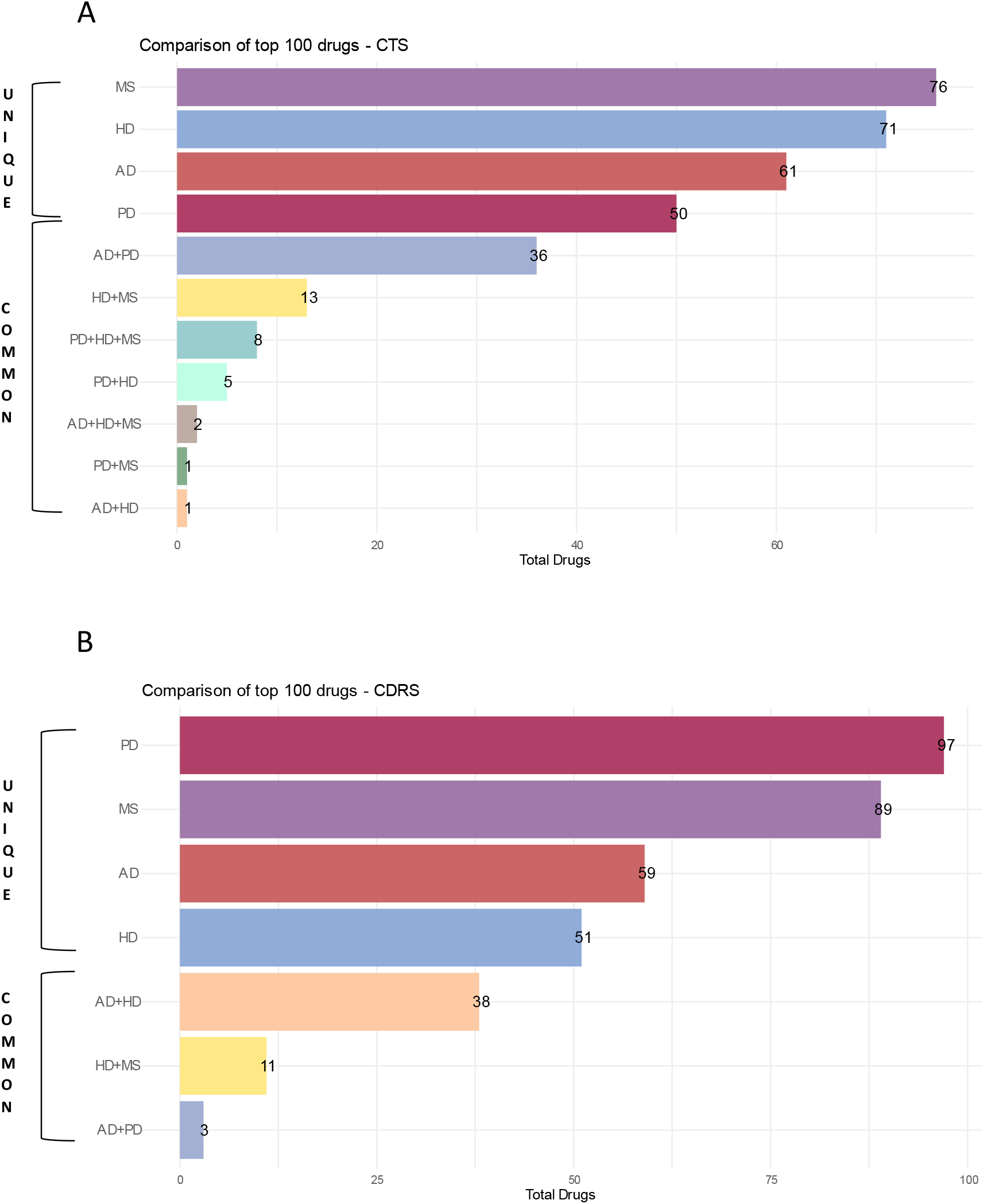
Comparison of the top 100 drugs generated from the super-reference table. **A**. Comparison of top 100 drugs using CTS and **B**. Comparison of top 100 drugs using CDRS. The top 100 drugs generated from the super-reference table of the Drug Repurposing Hub, for AD, PD, HD and MS were compared. Unique drugs for each disease are shown, as well as commonalities among diseases.

From the CTS Total Composite Scores, common drugs were detected in pairwise as in CDRS lists but also commonalities were detected among three diseases. Two common drugs were detected in AD, HD and MS (pipotiazine-palmitate, methylergometrine), while eight common drugs in PD, HD and MS (such as gyki-52466, ly215490, sym-2206 and talampanel). One common drug was detected in AD and HD, as well as in PD and MS (clozapine and lamotrigine respectively). Moreover, five common drugs were found in PD and HD such as budipine, amantadine and topiramate. Lastly, the highest number of commonalities was detected between AD and PD, with 36 common drugs, such as metergoline, tropisetron, zacopride and methysergide (Figure 9A).

### 3.5. Scoring new candidate drugs for repurposing against a selected disease

To present an application of the scoring part of D^R^e^A^mocracy, we scored different groups of drugs that are related to AD, such as the current FDA approved drugs, drugs that were previously failed in clinical trials for AD, as well as three different groups of candidate repurposed drugs that we suggested in our previous work for stage-specific drug repurposing for AD, for Braak stage I-II, III-IV and V-VI (Savva et al., 2022) (which was intentionally omitted from the sources used to compile the CDRS reference table).

Based on the output of D^R^e^A^mocracy, we compared the Composite Scores of the different drug groups that were tested. Figure 10 shows the comparison of the different drug groups, using the Composite scores of selected AD-related drugs (Braak I-II, Braak III-IV, Braak V-VI, FDA approved and Failed drugs) of the CDRS and CTS reference tables, as illustrated with boxplots (Panel A) and distribution histograms (Panel B). The Wilcoxon test was carried out to test whether the D^R^e^A^mocracy scores are significantly different across types of scores and groups of drugs of interest. Paired comparison of Braak I-II data using CDRS and CTS show statistical significance with a p-val=0.004. Moreover, paired comparison of Braak III-IV and Braak V-VI data using CTS and CDRS data, show statistical significance with a p-val=0.026 and 1.21e^-09^, respectively. When comparing the groups between them, statistical significance was found only for the comparison of Braak I-II versus Braak III-IV for the CTS reference table, with a p-val= 0.009.

**Figure 10:**
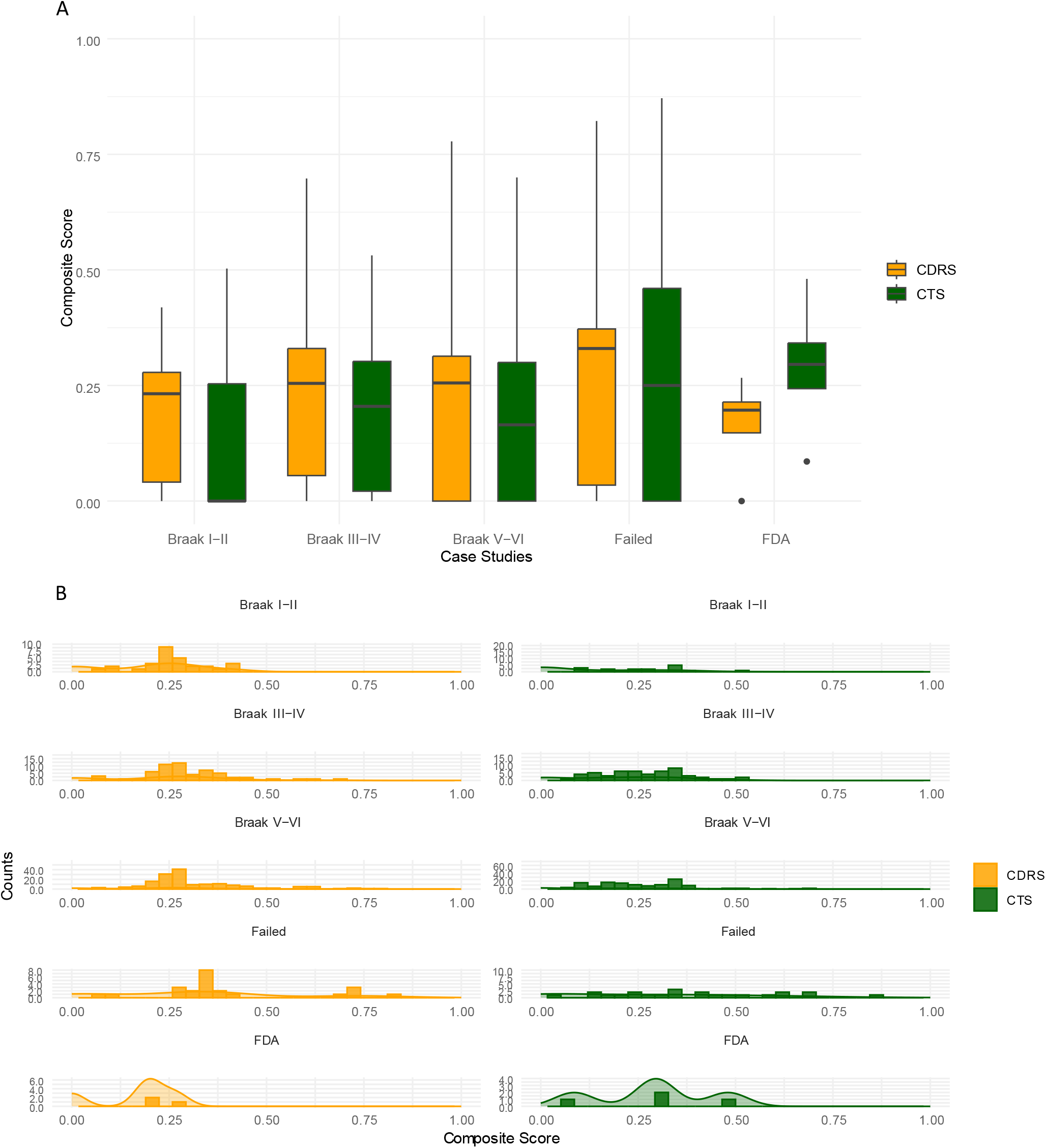
Plots of Composite scores for AD case examples. **A**. Box plots of Composite scores of selected AD-related drugs (Braak I-II, Braak III-IV, Braak V-VI, FDA approved and Failed) were generated using the CDRS and CTS reference tables. Black dots show the outliers. **B**. Distribution histogram of Composite Scores of selected AD-related drugs using the CDRS and CTS reference tables.

The output of D^R^e^A^mocracy in these test drug queries gives a snapshot of the trends regarding the selected drug categories and the existing knowledge from CTS and CDRS; FDA-approved drugs present higher Composite Scores as expected in CTS than in CDRS. CTS are expected to include candidate drugs with more similarity to the already FDA approved drugs whereas the candidate drugs from CDRS are expected to present higher diversity from the already approved drugs.

Additionally, failed drugs for AD have a higher median of Composite Score values in CDRS, than in CTS. Failed drugs could be supported more by the a-priori knowledge from CDRS than by the a-priori knowledge from CTS, as expected, since the diversity provided by the CDRS is higher than the one by CTS.

Among the three stage-specific groups of candidate repurposed drugs from our previous study [47], the candidate drugs for Braak stage I-II (incipient AD stage) are shown to be more unexpected based on the a priori knowledge from both CDRS and CTS collection methods. There is a noteworthy difference in the median values of the Scores between the two collection methods in the favor of CDRS as expected. Candidate drugs for Braak stage III-IV (moderate AD stage) and V-VI (severe AD stage) also present higher Composite Scores in CDRS than in CTS, suggesting that our computational repurposing study follows the trend of the a-priori knowledge of the rest of the CDRS as expected.

## 4. DISCUSSION

In this work we proposed a methodology, named D^R^e^A^mocracy, aiming to utilise a-priori knowledge to reveal trends in selected tracks of drug discovery research with increased resolution that includes modes of action, targeted pathways, initial indications for the investigated drugs, and to score new candidate drugs for repurposing against a selected disease. Four neurodegenerative diseases have been used as a testbed for the application of this methodology. Our analysis generated frequency tables, known as DC-matrices for each disease based on MoAs, Paths and initial Inds of the drugs detected from the computational drug repurposing studies and clinical trials. Based on these frequency tables, we constructed a super-reference table with drug suitability scores for each of the four neurodegenerative diseases. Hence, a given set of drugs can be scored based on the super-reference table with respect to its proposal in other studies (both clinical and computational repurposing ones). Moreover, disease to disease commonalities regarding Paths, MoAs and Inds were detected to highlight common mechanisms of neurodegeneration.

Following the creation of the super-reference table, we detected groups of top drugs for each disease based on the final total Composite score. From this analysis we detected common drugs at least in two diseases. For example, mianserin with a score of 0.8 for AD and a score of 0.7 for PD. Mianserin is a tetracyclic antidepressant and it strongly stimulates the release of norepinephrine [48]. Furthermore, ketanserin, gr-113808, gr125487, idalopirdine, r-1485, rs-23597-190, rs-39604, sb-203186, sb-258585, sb-271046, sb-399885 and sb-742457 with a score of 0.7 for AD and PD. Ketanserin is a selective serotonin receptor antagonist. Idalopirdine is a potent and selective 5-HT_6_ receptor which however has been discontinued for AD [49]. The majority of these drugs are serotonin inhibitors or antagonists.

In addition, the group of drugs iloperidone, pipotiazine-palmitate, clozapine, olanzapine, blonserin, paliperidone, lurasidone, zotepine, asepine, perospirone, quetiapine, ziprasidone and loxapine show high scores in three diseases, AD, PD and HD with a score of 0.7 for AD and 0.6 for both PD and HD. Iloperidone, clozapine, olanzapine and paliperidone are atypical antipsychotic for the treatment of schizophrenia symptoms. Iloperidone was also detected by our previous computational drug repurposing work [47] as the second highest scored candidate drug for Braak stage V-VI (severe) AD. A study by Choi et al. suggests a suppression of Aβ levels and plaque deposition in the brain of AD mice, as well as an improvement in the Aβ-induced memory impairment [50]. Additionally, clozapine therapy appeared to be beneficial in treatment-resistant agitation in patients with dementia [51]. Pipotiazine-palmitate is used for the maintenance treatment of chronic non-agitated schizophrenic patients [52]. All of these drugs are used in the treatment of psychotic and schizophrenia symptoms.

Moreover, we compared the output of our methodology with existing tools, such as DrugShot [2] and DrugEnrichr [53]. When comparing our results with DrugShot, a web-based server application that allows the extraction of ranked lists of drugs based on the search of a biomedical term, such as the disease of interest, we find both common and complementary outcomes. The top 200 drugs for AD based on the total Composite score and the top 200 drugs based on the publication count extracted from DrugShot were compared and we detected 18 common drugs between the two lists. Most of these drugs belong to the group of anti-depressive or anti-psychotic drugs, such as mirtazapine, aripiprazole, quetiapine, olanzapine, risperidone and others. Moreover, drugs such as amitriptyline, which is used for pain syndromes such as fibromyalgia, lamotrigine, which is used for epilepsy and bipolar disorder, celecoxib, a nonsteroidal anti-inflammatory drug, were detected as common between the two lists among others.

In our current study, four neurodegenerative diseases were used as case studies. For AD and PD, more data were available, specifically for CDRS, whereas for HD and MS, the two more under-represented diseases, less data were available regarding the CDRS collection method. This makes the use of D^R^e^A^mocracy more informative for AD and PD since more lists of repurposed drugs were available. The more drug lists available they are, the more accurate the methodology is. Additionally, D^R^e^A^mocracy leverages the existing knowledge to better understand where the research progress is currently focusing in both CTS and CDRS studies. Therefore, these findings do not include all possible mechanisms behind neurodegeneration, yet, it provides an overview of what most of the studies support at the current time being.

Our method gives the opportunity to researchers to compare their drugs based on prior knowledge from clinical trial studies and computational repurposed. What we can see from our findings and specifically from the frequency comparison between CDRS and CTS is that, most of the time, there is an analogous trend in the frequency between the two. These results suggest that the CDRS and CTS studies are consistent but not redundant, and hence complement each other. A combination of the two can enrich the individual information each collection method can give, and therefore, can lead to a more complete result. Nevertheless, it is very important to note that the addition of any other collection method such as experimental *in vitro* drug screens could further enhance the output of D^R^e^A^mocracy.

The limitations of the present study include the restriction of using a small, although curated, library of 6798 molecules, known as the Drug Repurposing Hub database. This means that drugs that need to be tested may not be included in the database and therefore, D^R^e^A^mocracy cannot score them. Moreover, drug names are used in their generic form, and hence a unique drug ID-based method given that this is supported by the required databases, should be used for the future. However, this study offers the ability to access information on what has been discovered so far. Future research should focus on the generation of other reference tables by collecting information on other diseases and also using the final total Composite score in combination with other scores such as the scoring scheme proposed in our previous work [47].

## 5. CONCLUSIONS

Overall, our study presents a novel methodology that takes advantage of the aggregated information from the drug repurposing lists already available as well as from clinical trial studies, towards the generation of a dynamic reference matrix with enhanced resolution regarding the disease-related frequencies of drug characteristics. We expect that our proposed methodology will be applied to many diseases of interest in the future and will serve as a reference for those interested in testing their drugs against the drug-related information available on these diseases.

## Data availability

The data used in this article are derived from sources in the public domain.

## Acknowledgement

None declared.

## Footnotes

1 C. P. Cavafy, “Ithaka” from C. P. Cavafy: Collected Poems. Translated by Edmund Keeley and Philip Sherrard. Translation Copyright © 1975, 1992 by Edmund Keeley and Philip Sherrard. Reproduced with permission of Princeton University Press. Source: C. P. Cavafy: Collected Poems (Princeton University Press, 1975)

